# Laparoscopic Roux-en-Y gastric bypass profoundly changes gut microbiota compared to laparoscopic sleeve gastrectomy: a metagenomic comparative analysis

**DOI:** 10.1101/425074

**Authors:** William Farin, Florian Plaza Oñate, Jonathan Plassais, Christophe Bonny, Christoph Beglinger, Bettina Woelnerhanssen, David Nocca, Frederic Magoules, Emmanuelle Le Chatelier, Nicolas Pons, Alessandra C.L. Cervino, S. Dusko Ehrlich

## Abstract

**Background:** Bariatric surgery is an effective therapeutic procedure for morbidly obese patients as it induces sustained weight loss. The two most common interventions are Laparoscopic Sleeve Gastrectomy (LSG) and Laparoscopic Roux-en-Y Gastric Bypass (LRYGB).

**Objective:** Characterizing the gut microbiota changes induced by LSG and LRYGB.

**Design:** 89 and 108 patients who underwent LSG and LRYGB respectively, were recruited from three countries: USA, France and Switzerland. Stools were collected before and 6 months after surgery. Microbial DNA was analysed with shotgun metagenomic sequencing (SOLiD 5500xl Wildfire). MSPminer, a novel innovative tool to characterize new in silico biological entities, was used to identify 715 Metagenomic Species Pan-genome (MSPs). 148 functional modules were analysed using GOmixer and KEGG database.

**Results:** Both interventions resulted in a similar increase of Shannon’s diversity index and gene richness of gut microbiota, in parallel with weight loss, but the changes of microbial composition were different. LRYGB led to higher relative abundance of aero-tolerant bacteria, such as *Escherichia coli* and buccal species, such as *Streptococcus* and *Veillonella* spp. In contrast, anaerobes such as *Clostridium* were more abundant after LSG, suggesting better conservation of anaerobic conditions in the gut. Function-level changes included higher potential for bacterial use of supplements such as vitamin B12, B1 and iron upon LRYGB. Moreover, after LRYGB, potential for nitrate and Trimethylamine oxidized (TMAO) respiration was detected.

**Conclusion:** Microbiota changes after bariatric surgery depend on the nature of the intervention. LRYGB induces greater taxonomic and functional changes in gut microbiota than LSG and may lead to a more dysbiotic microbiome. Possible long-term health consequences of these alterations remain to be established.

**Significance of this study:** *What is already known on this subject?:* Previous studies have reported changes of microbial composition after bariatric surgery with shotgun metagenomics, but lacked statistical power to document the changes. Important shifts in gut microbiome have been observed after LRYGB and LSG with an increase of aerotolerants from buccal microbiota. However, it is not clear how different the changes are between LRYGB and LSG although both procedures induce quite similar results with respect to weight loss, comorbidities remission and global glycaemic improvement outcomes.

*What are the new findings?:* LSG and LRYGB have specific but different impacts on gut microbial composition 6 months after bariatric surgery. The changes could be related to specific physiological effects. LRYGB promotes an important invasion of oral colonizers with *Veillonella* and *Streptococcus* genera. In contrast, these changes were less important after LSG. Moreover, microbial transportation potential of iron and vitamin B12 were also higher after LRYGB than LSG. We concluded that the type of surgery leads to different gut microbiome and functional profiles.

*How might it impact on clinical practice in the foreseeable future?:* Microbiome composition and functional profiles are not altered to the same extent by LRYGB and LSG 6 months after surgery. This difference should be considered when advising the patient on the type of bariatric surgery or post op diet.

## INTRODUCTION

The last decades have seen a dramatic increase in obesity rates worldwide. Nowadays obesity can be considered as a global epidemic. The prevalence of obesity in the USA keeps increasing and has reached 30% and Europe seems to follow a similar trend [1,2]. Obesity is associated to many other diseases such as type 2 diabetes (T2D), hyperlipidaemia, hypertension and obstructive sleep apnoea [3]. Bariatric surgery is currently the most effective strategy for morbidly obese patients with a sustainable weight loss and a success rate at five years higher than 66% [4]. The three main surgical procedures are laparoscopic Gastric Banding (LGB), Laparoscopic Roux-en-Y gastric bypass (LRYGB) and Laparoscopic Sleeve Gastrectomy (LSG) [5]. LRYGB consists in the division of the stomach into a small upper pouch and a larger remnant pouch by connecting the jejunum to the upper section. In that way, part of the stomach, the duodenum and the upper section of the jejunum are excluded from the passage. Alternatively, LSG is an irreversible procedure which does not imply bowel rearrangement, where a large portion of the stomach is removed along the great curvature. The last method (LGB) is the use of adjustable ring silicon-made around the upper stomach part to reduce food intake.

Nowadays, LRYGB and LSG are the two most often-used surgeries, as they appear to have better outcomes than LGB [5]. A recent report indicates no difference regarding the excessive BMI loss, quality of life, improvement of comorbidities and fatal consequences over a period of five years between LRYGB and LSG even if LRYGB was more efficient to treat gastrooesophageal reflux disease and dyslipidaemia [6].

Several studies support the hypothesis that gut microbiota plays a key role in obesity [7–10]. Energy metabolism, one of the major roles of gut microbiome, occurs through by-products such as short-chain fatty acids (SCFA) that result from microbial fermentation of carbohydrate fibres in the gut [11]. Individuals with lower gene richness are more prone to insulin resistance and dyslipidaemia [10]. *Akkermansia muciniphila*, has been consistently reported to be negatively associated to obesity and insulin sensitivity and its ability to degrade mucin can improve metabolic health during dietary interventions [12–14]. To investigate the impact of bariatric surgeries on gut microbiota, patients who have undergone either LRYGB or LSG were compared with their pre-operative (baseline) microbiota [15–21]. *A. muciniphila* and *E. coli* were both reported to be highly increased after LRYGB [15,16]. Liou *et al*., transplanted germfree mice with stool from LRYGB-treated mice and observed a reduction of fat mass, suggesting the gut composition induced by surgery has a direct effect on weight loss and host metabolism [16,22,23].

Metagenomics shotgun sequencing surpasses the limitations and biases of 16S rRNA based sequencing by providing higher resolution taxonomic and functional profiling [24]. To our knowledge, only a few studies used a shotgun sequencing based approach to characterize the microbiome modulation after LRYGB [15,21,22,25,26] and LSG [21,27,28]. However, previous reports [15,21,22,25–27] were limited by low number of patients, varying from n=6 to n=23, leading to a lack of statistical power. No direct comparison between the two interventions involving large cohort and their effect on the gut microbiome has been published.

Here, we present the largest metagenomics study to date investigating the effects of LRYGB and LSG on gut microbiome based on pre-operative and 6 month post-operative stools. This international multicentre study is based on 89 and 108 obese patients who underwent LRYGB and LSG respectively. In contrast to other studies relying on reference genomes, our analysis depends on a recent in silico method that discovers and quantifies microbial species including those hitherto unknown [29]. The goal is to understand taxonomic and functional changes that the two surgery types induce in the gut microbiome and also the differences between both interventions. Ultimately, a better understanding of the surgery effect can help to select the best type of surgery for the patients.

## MATERIAL AND METHODS

### Study population

In total, 275 patients diagnosed with obesity scheduled for LRYGB or LSG have been recruited in four clinical centres in three countries: France, Switzerland and USA. For the present investigation, inclusion criteria were age ≥18 years and Body Mass Index ≥ 35. Exclusion criteria were use of antibiotics and bowel cleansing for colonoscopy during the last two months before faecal sampling. The clinical protocol was approved by the institutional medical ethics committees.

### Data collection

Faecal samples were self-collected one month before and 6 months after the surgery and stored essentially as described by the International Human Microbiome Standards (IHMS) consortium standard operating protocol (SOP) 5 [30]. The medical records of participants were reviewed for demographic data, comorbidities and medications. In total, 531 faecal samples were collected before and 6 months after surgery from 275 patients. Patients with only one collected stool sample were excluded. In consequence, 197 patients were analysed with their respective 197 pre-operative and 197 post-operative stools that were collected and sequenced.

### DNA extraction and sequencing

Faecal DNA extraction was carried out according to the IHMS consortium SOP 7 [31]. Shotgun metagenomic sequencing was performed using SOLiD 5500xl Wildfire sequencing system (Life Technologies then ThermoFisher). 531 samples were sequenced yielding an average of 81.782 million (+/- 33.261 million) 35-base-long single reads.

### Reads mapping

Reads were cleaned using an in-house procedure included in METEOR software [32] in order to remove (i) those containing resilient sequencing adapters/barcodes and (ii) those with average quality less than 20. Cleaned reads were subsequently filtered from human and other possible food contaminant DNA (using Human genome RCh37-p10, *Bos taurus* and *Arabidopsis thaliana*) using Bowtie 1 [33] (2 mismatches permitted) included in METEOR software. The gene abundance profiling was based on the integrated catalogue of reference genes in the human gut microbiome [34]. Filtered high-quality reads were mapped to the 9.9M gene catalogue using Bowtie 1 (3 mismatches permitted) included in METEOR software. Using METEOR, the gene abundance profiling table was generated using a two-step procedure. First, the unique mapped reads (reads mapped to a unique gene in the catalogue) were attributed to their corresponding genes. Second, the shared reads (reads that mapped with the same alignment score to multiple genes in the catalogue) were attributed according to the ratio of their unique mapping counts. The counts were then normalized according to RPKM strategy (normalization by the gene size and the number of total mapped reads reported in frequency) to give the final gene relative abundance profile table.

### Taxonomic annotation

The genes were annotated using BLASTN alignment method against KEGG and RefSeq genomic databases [35,36]. The gene annotation method was adapted from Li et al. Only the hits with a minimum of 80% of query sequence length and 65% nucleotide identity were considered in the annotation process. The similarity thresholds for the phylum, genus and species taxonomic ranges were 65%, 80% and 95%, respectively. Genes with multiple hits deprived of any consensus (a consensus was defined as 10% of hits having the same annotation) for their taxonomic associations were annotated at a higher taxonomic range until a consensus was established.

### Functional annotation

Translated genes were aligned using BLASP version 2.6.0 against KEGG Release 81.0 (April 2017). Hits with a bitscore inferior to 60 or with an e-value superior to 0.01 were discarded. Each gene was finally assigned to the functional group (KEGG Ortholog or Enzyme Commission) associated with the most significant hit.

### Metagenomic Species Pan-genomes processing

The integrated reference catalogue of the human gut microbiome [34] was organized into 1696 Metagenomic Species Pan-genomes (MSPs) with MSPminer [29] by grouping co-abundant genes across 1267 stool samples coming from cohorts distinct of the one used in this study. For each MSP, the median vector of the square-root transformed counts of its core genes was computed by using the 531 samples of this study. MSPs detected in less than 5 samples were discarded. Then, the relationship between this median vector and the core genes was assessed with the Lin’s concordance correlation coefficient [37]. Finally, the abundance of the MSP was estimated from its 30 core genes with the highest concordance.

### Relative abundance estimation and feature selection

Out of 531 samples, 394 from 197 patients with two time-points were used for further analysis. The relative abundances at phylum level, based on the NCBI taxonomy, were computed by summing the relative abundances of all the genes belonging to the same phylum. The relative abundances of functional modules, which were based on the KEGG annotation, were computed by summing the relative abundances of all the genes belonging to the KEGG Orthologous (KO) groups. Abundances of 130 modules from GOmixer [38] were calculated by summing the abundances of each KO that belonged to the same module. This set of metabolic modules was selected because it was manually curated based on rigorous literature specific to gut bacterial functions. To increase important functional units that were lacking, 20 modules from KEGG were added to the GOmixer modules (**Supplementary Table 1**). The relative abundances of MSP were calculated by estimating the median of the 30 top genes belonging to its core genes. Microbial features that were present in <70 % of all the samples were removed. After filtering, statistical analysis was performed with the abundances of 15 phyla, 302 MSP and 5348 KEGG orthology (KO) groups. The values were log10 transformed, to approach normal distributions; a pseudo count equal to the lowest relative abundance value in the cohort was added to all relative abundances, to deal with zeros.

### Gene richness and taxonomic diversity

Gene richness for each sample was computed from the raw abundance table after downsizing based on 5 million simulated sequencing depth and 30 repeats. The gene richness was then re-calculated using a predictive model (linear regression) to be compared with a threshold of 11 million reads [10]. The Shannon index was computed to assess taxonomic diversity at both the genus and species level [39].

### Statistical test analysis

All statistical analyses were performed with R (version 3.3.2, http://www.R-project.org/). Clinical variables were summarized as medians with interquartile ranges (IQRs) or as frequencies with percentages. The Fisher’s exact test was used to compare categorical variables between patients who underwent LSG and LRYGB surgery.

### Statistical analysis to assess the effect of surgery on microbiota

First, we performed two independent analyses for LRYGB and for LSG. Regarding metagenomic variables, we used a non-parametric statistical test on log10 transformed relative abundances to show the microbial features that were significantly modulated by surgery. Since this is a prospective study and the samples are not independent, two-sided Wilcoxon signedrank test was used. Multiple testing was controlled by Benjamini-Hochberg (BH) false discovery rate (FDR). To ensure the MSP were significant, an additional filter based on the median fold change of relative abundance was applied. The median log2 fold change (FC) is the ratio of the median of this feature in all patients before surgery divided by the median of this feature in all patient after the gastric surgery. To summarize, a p-value lower than 0.05 and a (two-fold) FC >=1 or FC<=-1 were considered statistically significant.

### Statistical analysis to compare the two types of surgery

To evaluate the change of microbiome composition induced by the gastric surgery, the relative abundances from the post-operative stool were normalized by dividing them with relative abundances from pre-operative stools for each patient and then log2 transformed. The abundance ratios were then used to compare the surgery types by performing Wilcoxon rank test on the microbial features that were significant at least in one surgery group. Multiple testing was controlled by BH.

### Correlation between functional modules and MSPs

We performed two independent analyses for LRYGB and for LSG groups. In this context, Spearman correlation coefficients were calculated for every MSP that was found significantly impacted by gastric surgery and 150 functional modules. Relative abundances before and after surgery were used for the correlation and only correlation factors superior to 0.7 were retained for subsequent analysis.

## RESULTS

### Demographic description

197 patients have been included in three different countries (France, USA and Switzerland) before they underwent bariatric surgery (89 LRYGB and 108 LSG). Of these, 86 patients were included by two French clinical centres (Montpellier and Nice), 73 patients by Geisinger Health Center (USA) and 38 patients by St. Claraspital Basel (Switzerland). We checked if the presurgery characteristics of the cohort were homogeneous depending on the type of surgery performed (**Supplementary Table 2)**. Significant differences were found only for the country of origin (Fisher exact test, p<0.001) and diabetes prevalence (Fisher exact test, p<0.05). Important decrease of BMI was important after both surgical procedures (**Supplementary Table 3 and Table 4**).

### Analysis of gut microbiota before surgery

Since the surgery type was cofounded with the country of origin, we first analysed the differences in microbial gut composition at MSP level between the patients before surgery, using a Principal Component Analysis (PCA) of log-transformed relative abundances of MSPs (**Supplementary Figures 1.A and 1.B**). Patients were clearly separated by their country of origin and by the type of surgery performed. However, in the Swiss cohort, no difference was observed between the patients who underwent LRYGB (n=20) and LSG (n=18), (**Supplementary Figure 1.C**). To overcome this limitation, paired Wilcoxon rank test and ratio analysis were used to normalize the microbiome profile at baseline to reduce variability induced by country of origin.

### Surgery effects on gut microbial diversity and phylum-level composition

We estimated Shannon microbial diversity and gene richness in each sample. Compared to baseline, the Shannon index (**Supplementary Figure 2**) was significantly increased by surgery after 6 months (Wilcoxon signed-rank test, LRYGB: p=7.5*10^-6^ and LSG: p=7.3*10^-11^). Similarly, gene richness was increased very significantly (about 15%) by both procedures (LRYGB, p=3.8*10^-5^; LSG, p=5.9*10^-10^). As expected, 5 phyla were the most abundant in human intestinal gut: *Bacteroidetes*, *Firmicutes*, *Proteobacteria*, *Actinobacteria* and *Verrucomicrobia* (**Supplementary Figure 3**). Before surgery, phylum composition was comparable between the two groups. *Proteobacteria* was significantly increased by LRYGB (Wilcoxon signed-rank test, p=1.9*10^-12^) and LSG (Wilcoxon signed-rank test, p=5.4*10^-6^). *Firmicutes* showed a slight decrease after both surgeries.

### Surgery effects on microbiome species composition

51 MSP were significantly impacted by LRYGB (p<0.05 and FC_log2_ >=1 or FC_log2_ <=-1, Wilcoxon signed-rank tests) and 48 after LSG (**Figure 1A** and **Figure 2A; detailed view in Supplementary Figure 4 and 5, numerical values in supplementary information Table 5 and 6**). Of these, 20 were common to both procedures and were overwhelmingly (18 of 20) enriched upon surgery. Notably, the beneficial *Verrucomicrobia A. muciniphila* was enriched (LRYGB: FC_log2_=1.61, LSG: 1.82), but also the potentially proinflammatory *Proteobacteria* like *E. coli* (LRYGB: FC_log2_=5.22, LSG: FClog2=1.04), *Klebsiella pneumoniae* (LRYGB: FC_log2_=4.03, LSG: FClog2=2.09) and *Haemophilus parainfluenzae* (LRYGB: FC_log2_=1.61, LSG: FC_log2_=1.4). *Faecalibacterium prausnitzii* was less abundant only after LRYGB (FC_log2_=-1.43). Five oral species were enriched by both procedures, of which most strongly *Veillonela parvula* (LRYGB: FC_log2_=2.65, LSG: FC_log2_=1.92) and *Streptococcus salivarius* (LRYGB: FC_log2_=4.07, LSG: FC_log2_=1.98) but also *Streptococcus gordonii*, *Streptococcus mutans* and *Streptococcus parasanguinis*. Additional six oral species (*Fusobacteria nucleatum, Streptococcus anginosus, Streptococcus oralis, Streptococcus vestibularis, Veillonela atypica* and *Veillonela sp oral*) were enriched by LRYGB but not LSG. Interestingly, two oral *Bifidobacteria* were depleted by interventions, one by LRYGB (*Bifidobacteria bifidum*) and one by LSG (*Bifidobacteria dentium*). Four MSPs with no species-level annotation were enriched by both procedures (msp_0868, msp_0344, msp_0582 and msp_0355).

**Figure 1:**
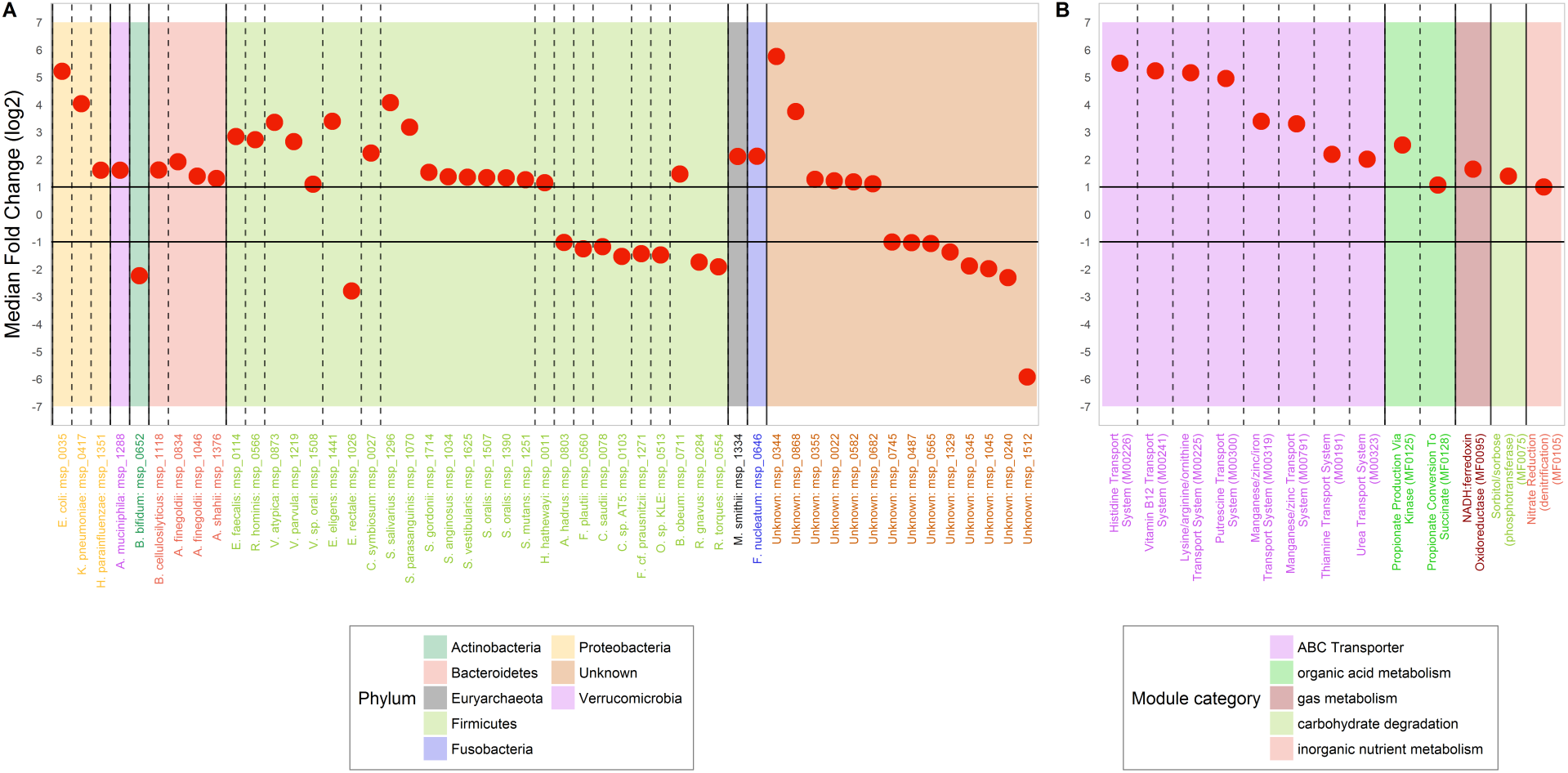
Microbial composition (A) and functional changes (B) following LRYGB.

**Figure 2:**
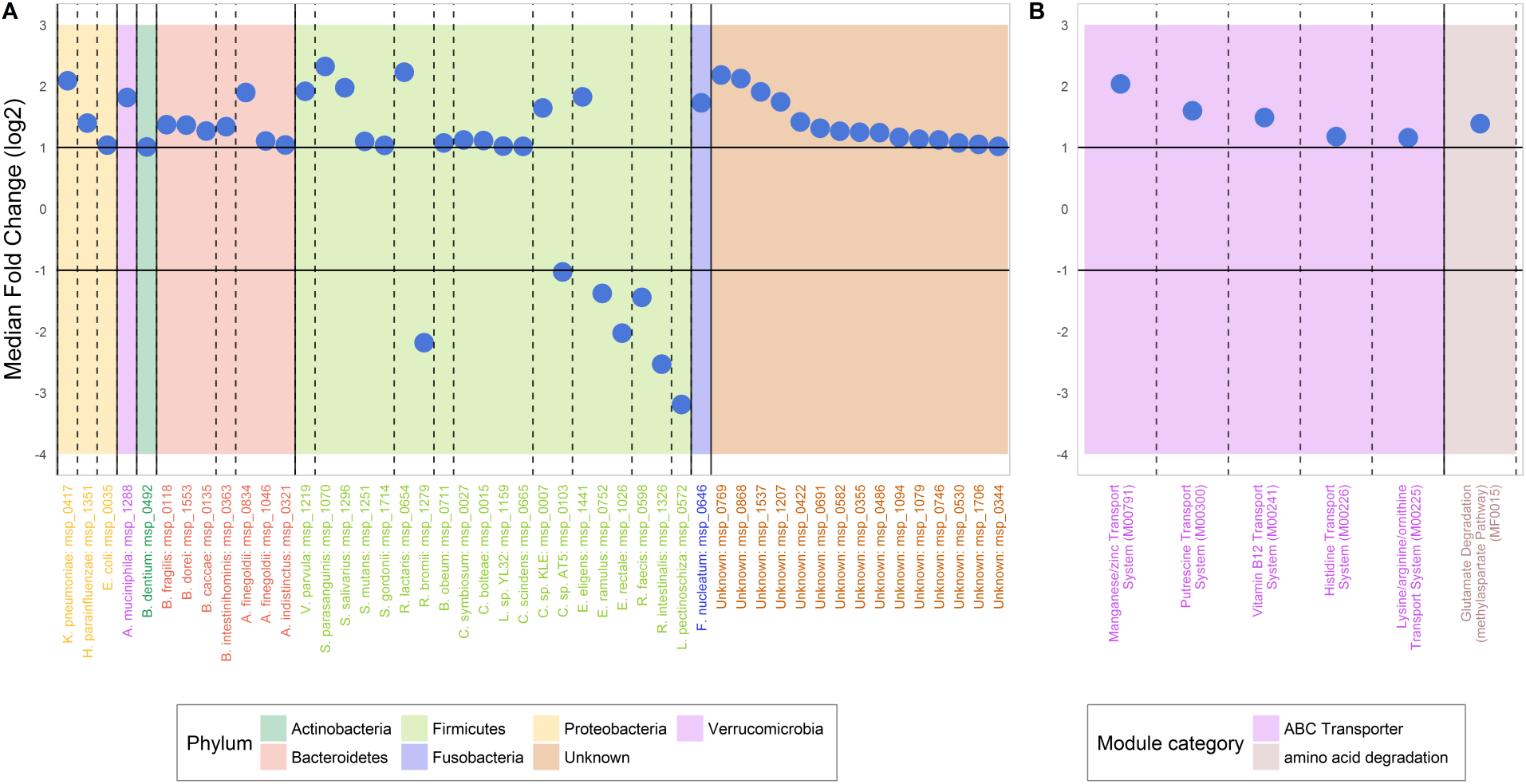
Microbial composition (A) and functional changes (B) following LSG.

Of the 78 MSPs impacted by LRYGB or/and LSG 24 had abundance ratios significantly different between LRYGB and LSG groups (p<0.05, Wilcoxon rank-sum tests) (**Figure 3A; Supplementary Table 7**). Two *Proteobacteria (E. coli* and *K. pneumonia*) and seven oral MSPs (two *S. oralis*, *S. parasanguinis, S. salivarius, V. atypica, V. parvula* and *V. sp. oral*) were enriched more by LRYGB than LSG surgery, one oral species, *B. bifidum* was more depleted. Two *Roseburia* (*R. faecis* and *R. hominis*) were also more enriched by LRYGB, as was *E. faecalis*. The five MSPs enriched more by LSG and annotated at the species level (*A. hadrus, C. sp KLE, F. plautii, O. sp. KLE* and *R. gnavus*) all belonged to *Firmicutes* order *Clostridiales*.

**Figure 3:**
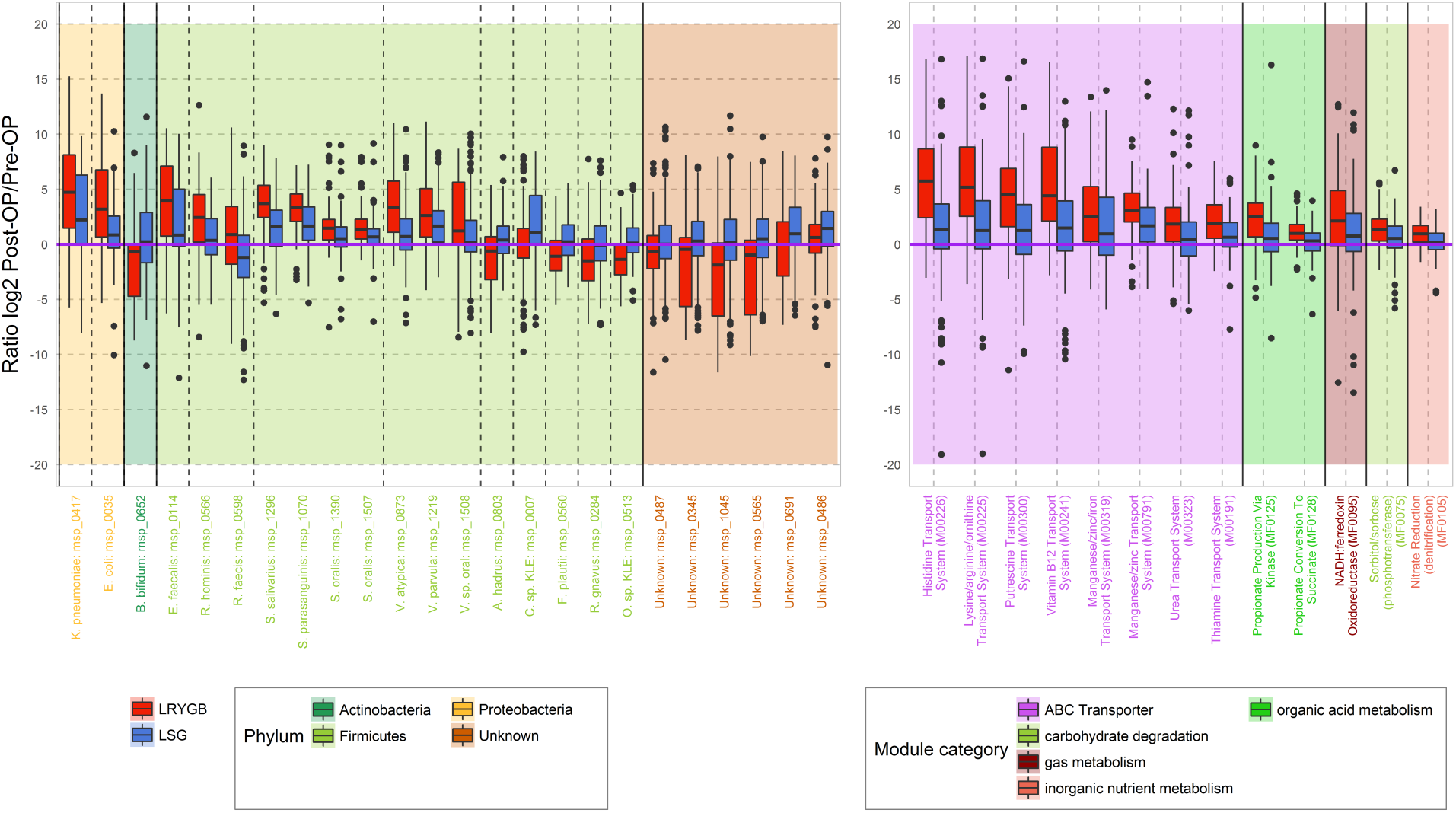
Differences of changes induced by LRYGB and LSG at species (A) and functionnal level (B).

### Effect of LRYGB and LSG on microbiome functions

13 functional modules were significantly more abundant after LRYGB and 6 after LSG; five were common to both interventions (**Figure 1B and Figure 2B, Supplementary information Table 8 and 9, Supplementary Figure 6 and 7**). Five common modules were ABC transporters implied in vitamin B12 (LRYGB: FC_log2_=5.23, LSG: FC_log2_=1.49), histidine (LRYGB: FC_log2_=5.51, LSG: FC_log2_=1.18), lysine/arginine (LRYGB: FC_log2_=5.16, LSG: FC_log2_=1.16), putrescin (LRYGB: FC_log2_=4.95, LSG: FC_log2_=1.6) and manganese/zinc (LRYGB: FC_log2_=3.30, LSG: FC_log2_=2.04) transport. Another manganese/zinc transport system was increased only after LYRGB (M00319; FC_log2_=3.39), as were Thiamine and Urea transport systems. Glutamate degradation module (FC_log2_=1.39) was more abundant after LSG while significant increase of nitrate reduction (FC_log2_=1.00) and propionate production (FC_log2_=2.53) modules were enriched after LRYGB.

Of the functional 14 modules affected by LRYGB or/and LSG 13 were found to be significantly different between LRYGB and LSG groups (**Figure 3B, Supplementary Table 10**). Notably, 8 functional modules involved in ABC transporters were more enriched in patients after LRYGB compared to the LSG surgery (histidine and lysine, putrescin, vitamin B12, manganese/zinc, urea and thiamine transport system). Modules involved in nitrate respiration and propionate production via kinase were more increased by LRYGB.

### Correlations of modules and MSPs

Spearman correlations between relative abundances of 78 MSP and 150 modules were computed independently for two groups of patients: LRYGB and LSG. Regarding the former, 8 modules were correlated to a set of five MSP annotated to *V. parvula*, *S. vestibularis*, *S. salivarius* and *S. parasanguinis* and *E. coli* (Spearman’s rho ≥ 0.7, **Supplementary Table 11**).

## DISCUSSION

### Microbiome composition is differentially altered by LRYGB and LSG

Physiological and anatomic changes of the gastrointestinal tract after bariatric surgery modify gut motility, gastric acid secretion, bile acid processing and gut hormone secretion [40]. Using whole metagenome shotgun sequencing with a large cohort of patients, we confirmed that gut microbiota was strongly modulated after LSG and LRYGB with notable similarities and differences. Gene richness was increased for most of the patients after both interventions. However, the most important divergence was the extent of the increase of *Proteobacteria* species. *E. coli and K. pneumoniae* were both increased by surgery but the increase was significantly stronger after LRYGB confirming results from other studies [15,25]. Increase of *E. coli* may reflect host and gut adaptation to maximize energy harvest in starvation-like conditions following bariatric surgery [18]. *A. muciniphila*, known to be negatively correlated to inflammation, was increased in patients after LSG or LRYGB in similar proportion in our study confirming results from other studies [15,25]. This species has been found to reverse obesity and increase mucus layer thickness in mice fed in HF-fed Diet [13]. These beneficial effects were summarized by the action of a membrane protein Amuc_1100 expressed by this bacterium [14]. Moreover, this increase was concomitant with a general improvement regarding the glycemic parameters after surgery.

In contrast, the number of MSP being negatively affected by bariatric surgery was lower. *F. prausnitzii*, which is a butyrate-producer, decreased 6 months after surgery in LRYGB while LSG had no effect on its presence in faeces. The decrease of *F. prausnitzii* in LYRGB patients was reported in two previous studies as well [15,25]. Similarly, *R. gnavus* and *R. torques* were also reportedly decreased in LRYGB. This species is well known to produce trans-scialidase to degrade mucin [41] and to be associated with inflammatory bowel diseases and metabolic disorders [10]. Some MSPs not annotated at species-level were also detected to be affected by both interventions. These observations were possible because MSPminer does not rely on reference genomes and can reveal new biological entities of interest.

### LRYGB promotes aerotolerant colonisation more than LSG

We observed that LRYGB led to a higher increase of oral colonizers (*Veillonela* and *Streptococcus* genus) than LSG. Possibly, less exposure to the acidic stomach compartment favours access of oral bacteria to the gut. However, facilitated access appears to be insufficient for implementation in the gut and other factors may be needed, as oral *Bifidobacteria* were depleted after surgery. Moreover, bypassing the duodenum might introduce some oxygen to the gastrointestinal tract [42], which is usually anaerobic, inhibiting growth of obligate anaerobes like *Clostridium* genus and promote domination of aerobes [43]. According to our results, *Clostridium* species were negatively impacted by LRYGB while they were enriched after LSG suggesting that the gut is still largely in anaerobic after LSG. Along the same lines, a higher relative abundance of ferredoxin oxidoreductase, which is usually associated with aerobic respiration, was observed after LRYGB relative to LSG. Evidence of oxidative stress was also reported with enrichment of functional modules involved in glutathione metabolism in LRYGB patients. For instance, biosynthesis glutathione was more increased in LRYGB (FC=0.76) than in LSG (FC=0.23), even if the FC_log2_ did not reach one in either case.

### Nitrates respiration may favour *E. coli* expansion after LRYGB

Following LRYGB, signs of nitrate reduction as an alternative form of respiration were observed. Interestingly, it has been recently observed that nitrate can boost growth of *E. coli* to outcompete species that rely on fermentation only [44,45]. LRYGB led also to enrichment of a functional module involved in urea transport system. This metabolite can be converted by urease to ammonia that can pass the mucosal barrier to reach the liver via portal vein. Bacterial urease is suspected to cause hyperammonemic encephalopathy in absence of liver cirrhosis [44]. Hyperammonemic encephalopathy has been recently described as being a severe complication of LRYGB [46]. After LRYGB, we also observed an increased potential for Trimethylamine oxidized (TMAO) utilization via pathways found in Proteobacteria (torYZ and torC, **Supplementary Table 12**). These results are in agreement with previous observations showing higher level of plasma circulating plasma TMAO levels in patients after LRYGB compared to LSG [47]. A retroconversion model of Trimethylamine (TMA) produced in gut by Enterobacteriaceae from TMAO reduction by gut bacteria has been reported [48].

### Microbial transportation of supplements is stimulated by bariatric surgery

We observed a strong increase of ABC transporters, especially Vitamin B12, B1 and manganese/iron transport systems after bariatric surgery. This was more pronounced in LRYGB, confirming previous findings [15]. After surgery, multivitamin, iron and calcium supplements are provided to compensate for deficiencies caused by food intake reduction and malabsorption [26]. Of note, in supplement tablets, vitamin B12, not bound to protein, is subsequently available to bacteria in the intestine. High levels of transport potential of vitamin B12, B1 and iron, particularly in LRYGB, suggest opportunist use of these nutrients by microbes. Transport of vitamin B12 was highly correlated with *E. coli* while iron transporter was associated with *S. salivarius* and *V. parvula* (**Supplementary Table 11**); these functions may have facilitated enrichment of the cognate species, acting synergistically with the decrease of exposure to acidic stomach compartment. High level of putrescin transport system was observed in post-operative stools after LRYGB and correlated with *E. coli* (**Supplementary Table 11**, Spearman’s rho = 0.85).

### Short chain fatty acids are altered by LRYGB and may impact weight loss via a GLP-1-dependant mechanism

SCFAs, metabolites formed by gut microbiota from carbohydrate substrates, can have several beneficial effects on the host. Butyrate and propionate were reported to be associated with weight loss and to have protective properties against diet-induced obesity in mice [49]. More precisely, the study indicated butyrate and propionate impact on gut hormones (by stimulating GLP-1 i.e glucagon-like peptide-1) and food intake reduction. In parallel, postprandial GLP-1 was observed to be significantly increased after LRYGB [50]. Abundance of propionate production module was higher after LRYGB and was also correlated with *E. coli* (Spearman’s rho = 0.76). In concordance to this observation, Ilhan has reported an increase of propionate after LRYGB [26]. Visceral and liver fat were also reduced after propionate delivery in humans [51]. General glycaemic improvement was more pronounced after LRYGB and could be related to higher gut propionate production. Breton et al. identified *ClpB*, a bacterial protein produced by *E. coli*, as an antigen-mimetic of a neuropeptide involved in the satietogenic system by stimulating GLP-1 [52]. The gene coding *ClpB* was found to be increased by 100 after LRYGB (p<0.05, Wilcoxon rank test; not shown), in parallel with the increase of *E. coli* that encodes it. Taken together, the data from literature and our results support that *E. coli* may potentially influence both host appetite and metabolism via a complex activation cascade.

## Conclusion

A direct comparison of microbiome changes after LSG and LRYGB procedures, assessed via large cohorts and shotgun metagenomic indicated a profound modification of bacterial gut composition 6 months after bariatric surgery in parallel with weight loss. Dramatic increase of aero-tolerant bacteria from phylum *Proteobacteria* suggests physiological changes that are specific to LRYGB; the presence of potentially pro-inflammatory bacteria in the microbiome may sustain low-level inflammation and thus impact evolution of the disease. Long-terms observations from surgery, possibly 3-5 years, are necessary to highlight the clinical relevance of these findings. We infer from these data that LRYGB had greater impact on microbiome composition than LSG, leading to a potential gut dysbiosis. This could be considered when advising the patient on the type of bariatric surgery or post op diet.

## ACKNOWLEDGEMENTS

We thank Prof. Ralph Peterli to have performed the operations and recruiting for the Swiss cohort and Dr Anne Christin Meyer-Gerspach who was very helpful to collect all stool samples. We thank Rodolphe Anty to have recruit patients from CHU Nice. We also thank Karine Roger and Rachel Morra to have monitored the clinical study and regrouped the clinical data. We finally thank Pierre Rimbaud for his thoughtful comments and suggestions to improve the manuscript.

## Abbreviations

BH: Benjamini-Hochberg
FDR: False Discovery Rate
IHMS: International Human Microbiome Standards
IQR: interquartile ranges
LGB: Laparoscopic Gastric Banding
LRYGB: Laparoscopic Roux-en-Y Gastric Bypass
LSG: Laparoscopic Sleeve Gastrectomy
MSP: Metagenomic Species Pan-genomes
SCFA: Short-Chain Fatty Acids
SOP: Standard Operating Protocol
TMA: Trimethylamine
TMAO: Trimethylamine oxidized

## References

1 Hruby A, Hu FB. The Epidemiology of Obesity: A Big Picture. PharmacoEconomics 2015;33:673–89. doi:10.1007/s40273-014-0243-x

2 Gallus S, Lugo A, Murisic B, et al. Overweight and obesity in 16 European countries. Eur J Nutr 2015;54:679–89. doi:10.1007/s00394-014-0746-4

3 Jarolimova J, Tagoni J, Stern TA. Obesity: Its Epidemiology, Comorbidities, and Management. Prim Care Companion CNS Disord 2013;15. doi:10.4088/PCC.12f01475

4 Nocca D, Loureiro M, Skalli EM, et al. Five-year results of laparoscopic sleeve gastrectomy for the treatment of severe obesity. Surg Endosc 2017;31:3251–7. doi:10.1007/s00464-016-5355-2

5 Colquitt JL, Pickett K, Loveman E, et al. Surgery for weight loss in adults. Cochrane Database Syst Rev 2014;:CD003641. doi:10.1002/14651858.CD003641.pub4

6 Peterli R, Wölnerhanssen BK, Peters T, et al. Effect of Laparoscopic Sleeve Gastrectomy vs Laparoscopic Roux-en-Y Gastric Bypass on Weight Loss in Patients With Morbid Obesity: The SM-BOSS Randomized Clinical Trial. JAMA 2018;319:255–65. doi:10.1001/jama.2017.20897

7 Baothman OA, Zamzami MA, Taher I, et al. The role of Gut Microbiota in the development of obesity and Diabetes. Lipids Health Dis 2016;15:108. doi:10.1186/s12944-016-0278-4

8 Turnbaugh PJ, Ley RE, Mahowald MA, et al. An obesity-associated gut microbiome with increased capacity for energy harvest. Nature 2006;444:1027–31. doi:10.1038/nature05414

9 Turnbaugh PJ, Gordon JI. The core gut microbiome, energy balance and obesity. J Physiol 2009;587:4153–8. doi:10.1113/jphysiol.2009.174136

10 Le Chatelier E, Nielsen T, Qin J, et al. Richness of human gut microbiome correlates with metabolic markers. Nature 2013;500:541–6. doi:10.1038/nature12506

11 Morrison DJ, Preston T. Formation of short chain fatty acids by the gut microbiota and their impact on human metabolism. Gut Microbes 2016;7:189–200. doi:10.1080/19490976.2015.1134082

12 Dao MC, Everard A, Aron-Wisnewsky J, et al. Akkermansia muciniphila and improved metabolic health during a dietary intervention in obesity: relationship with gut microbiome richness and ecology. Gut 2016;65:426–36. doi:10.1136/gutjnl-2014-308778

13 Everard A, Belzer C, Geurts L, et al. Cross-talk between Akkermansia muciniphila and intestinal epithelium controls diet-induced obesity. Proc Natl Acad Sci U S A 2013;110:9066–71. doi:10.1073/pnas.1219451110

14 Plovier H, Everard A, Druart C, et al. A purified membrane protein from Akkermansia muciniphila or the pasteurized bacterium improves metabolism in obese and diabetic mice. Nat Med 2017;23:107–13. doi:10.1038/nm.4236

15 Palleja A, Kashani A, Allin KH, et al. Roux-en-Y gastric bypass surgery of morbidly obese patients induces swift and persistent changes of the individual gut microbiota. Genome Med 2016;8:67. doi:10.1186/s13073-016-0312-1

16 Liou AP, Paziuk M, Luevano J-M, et al. Conserved Shifts in the Gut Microbiota Due to Gastric Bypass Reduce Host Weight and Adiposity. Sci Transl Med 2013;5:178ra41. doi:10.1126/scitranslmed.3005687

17 Kong L-C, Tap J, Aron-Wisnewsky J, et al. Gut microbiota after gastric bypass in human obesity: increased richness and associations of bacterial genera with adipose tissue genes. Am J Clin Nutr 2013;98:16–24. doi:10.3945/ajcn.113.058743

18 Furet J-P, Kong L-C, Tap J, et al. Differential adaptation of human gut microbiota to bariatric surgery-induced weight loss: links with metabolic and low-grade inflammation markers. Diabetes 2010;59:3049–57. doi:10.2337/db10-0253

19 Zhang H, DiBaise JK, Zuccolo A, et al. Human gut microbiota in obesity and after gastric bypass. Proc Natl Acad Sci 2009;106:2365–70. doi:10.1073/pnas.0812600106

20 Medina DA, Pedreros JP, Turiel D, et al. Distinct patterns in the gut microbiota after surgical or medical therapy in obese patients. PeerJ 2017;5:e3443. doi:10.7717/peerj.3443

21 Murphy R, Tsai P, Jüllig M, et al. Differential Changes in Gut Microbiota After Gastric Bypass and Sleeve Gastrectomy Bariatric Surgery Vary According to Diabetes Remission. Obes Surg 2017;27:917–25. doi:10.1007/s11695-016-2399-2

22 Tremaroli V, Karlsson F, Werling M, et al. Roux-en-Y Gastric Bypass and Vertical Banded Gastroplasty Induce Long-Term Changes on the Human Gut Microbiome Contributing to Fat Mass Regulation. Cell Metab 2015;22:228–38. doi:10.1016/j.cmet.2015.07.009

23 Li JV, Ashrafian H, Bueter M, et al. Metabolic surgery profoundly influences gut microbial-host metabolic cross-talk. Gut 2011;60:1214–23. doi:10.1136/gut.2010.234708

24 Ranjan R, Rani A, Metwally A, et al. Analysis of the microbiome: Advantages of whole genome shotgun versus 16S amplicon sequencing. Biochem Biophys Res Commun 2016;469:967–77. doi:10.1016/j.bbrc.2015.12.083

25 Graessler J, Qin Y, Zhong H, et al. Metagenomic sequencing of the human gut microbiome before and after bariatric surgery in obese patients with type 2 diabetes: correlation with inflammatory and metabolic parameters. Pharmacogenomics J 2013;13:514–22. doi:10.1038/tpj.2012.43

26 Ilhan ZE, DiBaise JK, Isern NG, et al. Distinctive microbiomes and metabolites linked with weight loss after gastric bypass, but not gastric banding. ISME J 2017;11:2047–58. doi:10.1038/ismej.2017.71

27 Damms-Machado A, Mitra S, Schollenberger AE, et al. Effects of surgical and dietary weight loss therapy for obesity on gut microbiota composition and nutrient absorption. BioMed Res Int 2015;2015:806248. doi:10.1155/2015/806248

28 Liu R, Hong J, Xu X, et al. Gut microbiome and serum metabolome alterations in obesity and after weight-loss intervention. Nat Med 2017;23:859–68. doi:10.1038/nm.4358

29 Oñate FP, Cervino ACL, Magoulès F, et al. Abundance-based reconstitution of microbial pangenomes from whole-metagenome shotgun sequencing data. bioRxiv 2017;:173203. doi:10.1101/173203

30 SOPs05. http://www.microbiome-standards.org/index.php?id=292 (accessed 9 Jan 2018).

31 SOPs07. http://www.microbiome-standards.org/index.php?id=254) (accessed 9 Jan 2018).

32 Pons N, Batto J-M, Kennedy S, et al. JOBIM 2010 NPONS POSTER. 2014.

33 Langmead B, Trapnell C, Pop M, et al. Ultrafast and memory-efficient alignment of short DNA sequences to the human genome. Genome Biol 2009;10:R25. doi:10.1186/gb-2009-10-3-r25

34 Li J, Jia H, Cai X, et al. An integrated catalog of reference genes in the human gut microbiome. Nat Biotechnol 2014;32:834–41. doi:10.1038/nbt.2942

35 Kanehisa M, Goto S, Sato Y, et al. Data, information, knowledge and principle: back to metabolism in KEGG. Nucleic Acids Res 2014;42:D199–205. doi:10.1093/nar/gkt1076

36 Tatusova T, Ciufo S, Fedorov B, et al. RefSeq microbial genomes database: new representation and annotation strategy. Nucleic Acids Res 2014;42:D553-559. doi:10.1093/nar/gkt1274

37 Lin LI. A concordance correlation coefficient to evaluate reproducibility. Biometrics 1989;45:255–68.

38 Darzi Y, Falony G, Vieira-Silva S, et al. Towards biome-specific analysis of meta-omics data. ISME J 2016;10:1025–8. doi:10.1038/ismej.2015.188

39 BSTJ 28: 4. October 1949: Communication Theory of Secrecy Systems. (Shannon, C.E.). 1949. http://archive.org/details/bstj28-4-656 (accessed 9 Jan 2018).

40 Quercia I, Dutia R, Kotler DP, et al. Gastrointestinal changes after bariatric surgery. Diabetes Metab 2014;40:87–94. doi:10.1016/j.diabet.2013.11.003

41 Crost EH, Tailford LE, Monestier M, et al. The mucin-degradation strategy of Ruminococcus gnavus: The importance of intramolecular trans-sialidases. Gut Microbes 2016;7:302–12. doi:10.1080/19490976.2016.1186334

42 Celiker H. A new proposed mechanism of action for gastric bypass surgery: Air hypothesis. Med Hypotheses 2017;107:81–9. doi:10.1016/j.mehy.2017.08.012

43 Hartman AL, Lough DM, Barupal DK, et al. Human gut microbiome adopts an alternative state following small bowel transplantation. Proc Natl Acad Sci 2009;106:17187–92. doi:10.1073/pnas.0904847106

44 Tiso M, Schechter AN. Nitrate Reduction to Nitrite, Nitric Oxide and Ammonia by Gut Bacteria under Physiological Conditions. PLoS ONE 2015;10. doi:10.1371/journal.pone.0119712

45 Winter SE, Winter MG, Xavier MN, et al. Host-derived nitrate boosts growth of E. coli in the inflamed gut. Science 2013;339:708–11. doi:10.1126/science.1232467

46 Nagarur A, Fenves AZ. Late presentation of fatal hyperammonemic encephalopathy after Roux-en-Y gastric bypass. Proc Bayl Univ Med Cent 2017;30:41–3.

47 Schultes B. Increased Trimethylamine-N-Oxide (TMAO) Levels After Roux-en Y Gastric Bypass Surgery-Should We Worry About It? Obes Surg 2017;27:2170–3. doi:10.1007/s11695-017-2757-8

48 Hoyles L, Jiménez-Pranteda ML, Chilloux J, et al. Metabolic retroconversion of trimethylamine N-oxide and the gut microbiota. bioRxiv 2017;:225581. doi:10.1101/225581

49 Lin HV, Frassetto A, Kowalik Jr EJ, et al. Butyrate and Propionate Protect against Diet-Induced Obesity and Regulate Gut Hormones via Free Fatty Acid Receptor 3-Independent Mechanisms. PLoS ONE 2012;7. doi:10.1371/journal.pone.0035240

50 Jirapinyo P, Jin DX, Qazi T, et al. A Meta-Analysis of GLP-1 After Roux-En-Y Gastric Bypass: Impact of Surgical Technique and Measurement Strategy. Obes Surg 2018;28:615–26. doi:10.1007/s11695-017-2913-1

51 Chambers ES, Viardot A, Psichas A, et al. Effects of targeted delivery of propionate to the human colon on appetite regulation, body weight maintenance and adiposity in overweight adults. Gut 2015;64:1744–54. doi:10.1136/gutjnl-2014-307913

52 Breton J, Tennoune N, Lucas N, et al. Gut Commensal E. coli Proteins Activate Host Satiety Pathways following Nutrient-Induced Bacterial Growth. Cell Metab 2016;23:324–34. doi:10.1016/j.cmet.2015.10.017

